# A sexually dimorphic neuroendocrine cascade acts through shared sensory neurons to regulate behavior in *C. elegans*

**DOI:** 10.1101/281741

**Authors:** Zoë A. Hilbert, Dennis H. Kim

## Abstract

**ABSTRACT**

Sexually dimorphic behaviors are observed in species across the animal kingdom, however the relative contributions of sex-specific and sex-shared nervous systems to such behaviors are not fully understood. Building on our previous work which described the sexually dimorphic expression of a neuroendocrine ligand, DAF-7, and its role in behavioral decision-making in *C. elegans* (Hilbert and Kim, 2017), we show here that sex-specific expression of *daf-7* is regulated by another neuroendocrine ligand, Pigment Dispersing Factor (PDF-1), which has previously been implicated in regulating male-specific behavior (Barrios et al., 2012). Our analysis revealed that PDF-1 acts sex- and cell-specifically in the ASJ neurons to regulate the expression of *daf-7* and we show that differences in the expression of the PDFR-1 receptor account for the sex-specific effects of this pathway. Our data suggest that modulation of the sex-shared nervous system by neuroendocrine signaling pathways can play a role in shaping sexually dimorphic behaviors.

## INTRODUCTION

Behavioral differences between the sexes of many animal species can make major contributions to the reproductive fitness of the organism. While sex-specific behaviors can be readily observed, the mechanistic basis of such behavioral differences is less well understood. Morphological differences, including the existence of sex-specific neurons, have been documented in the nervous systems of many species, but differences in sex-shared neurons have also been implicated in generating sex-specific behaviors. In particular, how sex-specific behavioral circuits are generated within the features of the nervous system common to both sexes has been the focus of recent studies in diverse organisms. Work on the mouse vomeronasal organ (VNO) has suggested that the functional circuits for both male- and female-specific behaviors such as courtship and aggression are intact in the brains of both sexes and are modulated by VNO activity in response to pheromone cues (Kimchi et al., 2007; Stowers et al., 2002). In a similar vein, the *Drosophila* male pheromone 11-*cis* Vaccenyl acetate (cVa) has been shown to be sensed by the same neurons in the two sexes but stimulates distinct sex-specific behavioral responses (Datta et al., 2008; Kohl et al., 2013; Kurtovic et al., 2007; Ruta et al., 2010). These examples and others have provided some insight into sexual dimorphisms in the nervous system although many open questions remain (Dulac and Kimchi, 2007; Stowers and Logan, 2010; Yang and Shah, 2014).

In the nematode *Caenorhabditis elegans*, behavioral differences between the two sexes— hermaphrodites and males—range from behaviors exclusively performed by one sex, such as egg laying by hermaphrodites and the unique mating program of males (Liu and Sternberg, 1995), to those in which the two sexes differ in their responses to the same stimuli, including differing responses to pheromone (Fagan et al., 2018; Jang et al., 2012; Srinivasan et al., 2008), food-related cues (Ryan et al., 2014), and conditioning to aversive stimuli (Sakai et al., 2013; Sammut et al., 2015). While sex-specific neurons regulate corresponding behaviors in *C. elegans*, the 294 neurons that are common to the nervous systems of both hermaphrodites and males have emerged as major contributors to a number of different sexually dimorphic behaviors (Barr et al., 2018; Barrios et al., 2012; 2008; Fagan et al., 2018; Lee and Portman, 2007; Mowrey et al., 2014; Sakai et al., 2013). In particular, recent work has uncovered sexually dimorphic differences in axonic and dendritic morphology and synaptic connectivity within the sex shared nervous system, which can modulate neuronal circuits and behavior (Hart and Hobert, 2018; Oren-Suissa et al., 2016; Serrano-Saiz et al., 2017a; Weinberg et al., 2018). In addition, studies of sexually dimorphic gene expression (Hilbert and Kim, 2017; Ryan et al., 2014; Serrano-Saiz et al., 2017a) and neurotransmitter identity (Gendrel et al., 2016; Pereira et al., 2015; Serrano-Saiz et al., 2017a; 2017b) have suggested that sexual differentiation of neurons within the sex-shared nervous system of *C. elegans* is also critical for the establishment of sexually dimorphic behaviors.

We have previously demonstrated that *daf-7*, which encodes a TGF*β* family neuroendocrine ligand that regulates diverse aspects of *C. elegans* behavior and physiology (Chang et al., 2006; Fletcher and Kim, 2017; Gallagher et al., 2013; Greer et al., 2008; Milward et al., 2011; Ren et al., 1996; Shaw et al., 2007; White and Jorgensen, 2012; You et al., 2008), is expressed in a sex-specific and context-dependent manner in the sex-shared ASJ chemosensory neurons and functions to promote exploratory behaviors (Hilbert and Kim, 2017; Meisel et al., 2014). Regulation of *daf-7* expression in the ASJ neurons requires the integration of sensory and internal state information including the sex and age of the animal, its nutritional state, and the type of bacterial species it encounters in its environment (Hilbert and Kim, 2017). These stimuli feed into the regulation of *daf-7* expression in the two ASJ neurons in a hierarchical manner, which enables the animal to make behavioral decisions taking into account past experiences as well as its current environment.

Here we report the identification of a second neuroendocrine signaling pathway, the Pigment Dispersing Factor (PDF-1) pathway, which functions to regulate the expression of *daf-7* and its effects on behavior in a sex-specific manner. We show that PDF-1 pathway signaling, which has previously been shown to be essential for male mate-searching behavior (Barrios et al., 2012), functions sex-specifically in the ASJ neurons themselves to regulate *daf-7* expression. Further, we demonstrate that the sex-specificity of PDF-1 regulation of *daf-7* derives from differences in the expression of the PDF-1 receptor gene, *pdfr-1*, in the ASJ neurons. Our data suggest that the gating of neuronal responses to neuropeptide modulators through sex-specific restriction of receptor expression is a mechanism by which sex-specific behaviors can be generated from the largely sex-shared nervous system in *C. elegans*.

## RESULTS AND DISCUSSION

### PDF-1 neuropeptide signaling regulates the sex-specific expression of *daf-7* in the ASJ chemosensory neurons

To explore the molecular and genetic mechanisms that underlie the sex-specificity of *daf-7* expression, we identified a number of candidate genes that had previously been shown to be involved in the regulation of mate-searching behavior or other aspects of male physiology and tested mutants of these genes for effects on *daf-7* expression in the male ASJ neurons. Through this approach, we identified the PDF-1 neuropeptide signaling pathway as a regulator of *daf-7* expression in the ASJ neurons (Figure 1). The PDF neuropeptide signaling pathway is conserved among insects, crustaceans and nematodes. In *Drosophila melanogaster*, PDF signaling has been well studied for its critical role in the regulation of circadian rhythmicity (Helfrich-Förster, 1995; Park and Hall, 1998; Park et al., 2000; Renn et al., 1999), but it has also been shown to modulate geotaxis (Mertens et al., 2005), pheromone production and mating behaviors (Fujii and Amrein, 2010; Kim et al., 2013; Krupp et al., 2013). In *C. elegans*, PDF-1 signaling has been established as an important regulator of locomotion, roaming behaviors, quiescence, and notably, male mate-searching behavior (Figure 1A; Barrios et al., 2012; Choi et al., 2013; Flavell et al., 2013; Janssen et al., 2008; 2009; Meelkop et al., 2012).

**Figure 1.**
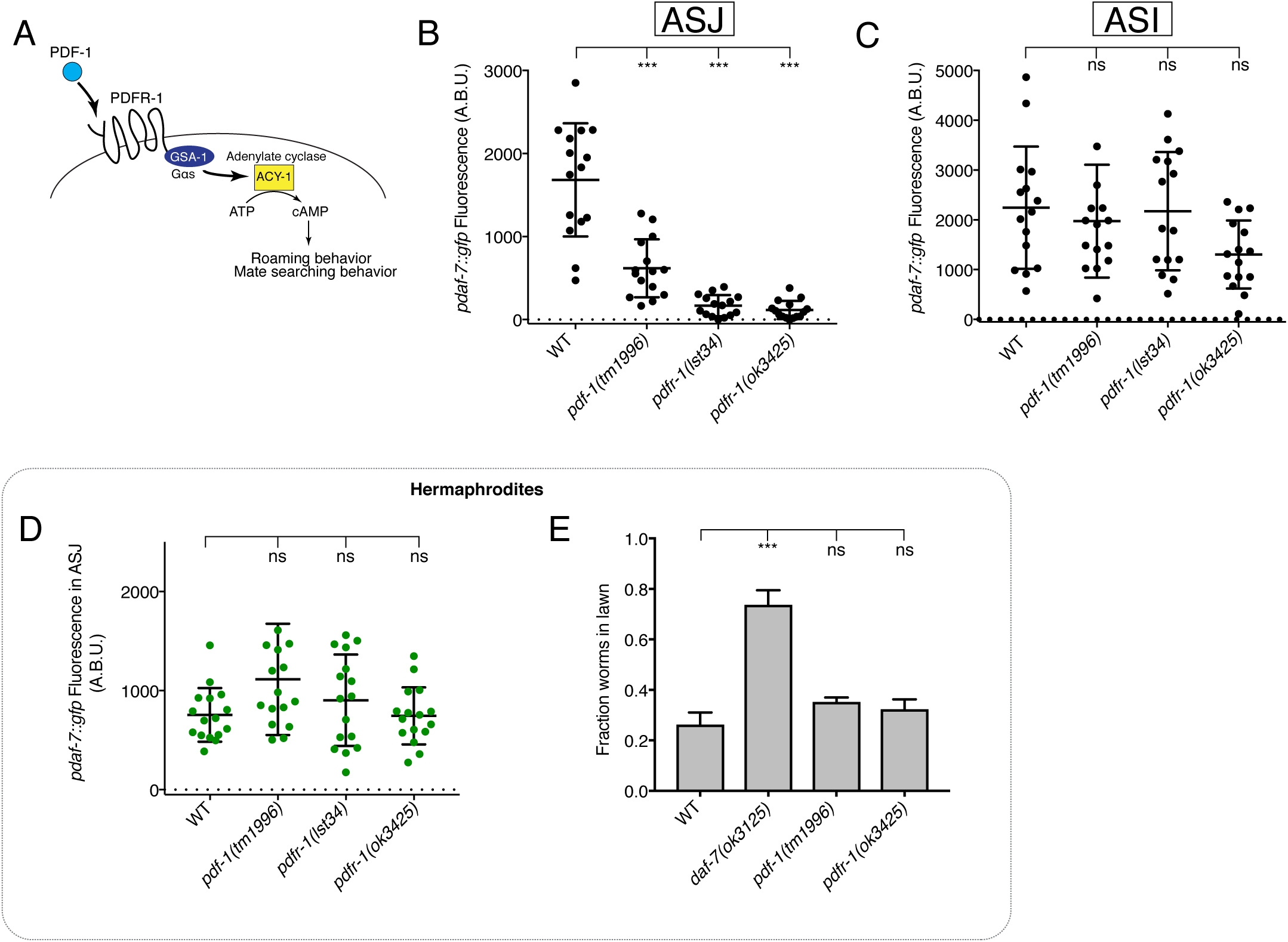
The PDF-1 pathway is required for the male-specific expression of *daf-7/*TGF*β* in the ASJ neurons. (A) PDF-1 signaling activates cAMP production and regulates both roaming behavior and male mate-searching behavior in *C. elegans*. (B-C) Maximum fluorescence values of *pdaf-7::gfp* in the ASJ (B) and ASI (C) neurons of adult male animals. ***p< 0.001 as determined by ordinary one-way ANOVA followed by Dunnett’s multiple comparisons test. Error bars represent standard deviation (SD). ns, not significant. n=15 animals for all genotypes. (D) Maximum fluorescence values of *pdaf-7::gfp* in the ASJ neurons of hermaphrodites after 16 hours on *P. aeruginosa*. Significance determined by ordinary one-way ANOVA followed by Dunnett’s multiple comparisons test. Error bars represent SD. ns, not significant. n=15 animals for all genotypes. (E) Lawn occupancy of animals on *P. aeruginosa* after 24 hours. ***p< 0.001 as determined by ordinary one-way ANOVA followed by Dunnett’s multiple comparisons test. Values plotted indicate the mean + SD for three replicates. Number of animals assayed are as follows: WT (n=89), *daf-7* (n=66), *pdf-1* (n=117), *pdfr-1* (n=105).

We observed that males with mutation of either the PDF-1 neuropeptide ligand or its receptor, PDFR-1, have markedly attenuated expression of *daf-7* in the ASJ neuron pair (Figure 1B). The ASI chemosensory neurons are established sites of *daf-7* expression in both male and hermaphrodite animals (Ren et al., 1996; Schackwitz et al., 1996), so we asked if the PDF-1 signaling pathway also regulates *daf-7* expression in these neurons. In the PDF-1 pathway mutant males, we observe no difference in *daf-7* levels in the ASI neurons when compared to WT (Figure 1C), suggesting that the PDF-1 pathway specifically affects the regulation of *daf-7* in the ASJ neuron pair.

We have previously reported that *daf-7* expression serves a dual role in the ASJ neurons, functioning in males to promote food-leaving behaviors (Hilbert and Kim, 2017), but also being induced by the presence of pathogenic bacteria such as *Pseudomonas aeruginosa* in both sexes to promote pathogen avoidance behaviors (Meisel et al., 2014). Given this and our interest in identifying male-specific regulators of *daf-7* expression, we asked if the PDF-1 pathway is required for the upregulation of *daf-7* expression in response to *P. aeruginosa* exposure. We did not observe a requirement for PDF-1 signaling in the induction of *daf-7* expression in the ASJ neurons after 16 hours on *P. aeruginosa*; both *pdf-1* and *pdfr-1* mutant hermaphrodites had equivalent levels of *daf-7* expression when compared to control animals (Figure 1D). Similarly, males that are mutant for either the PDF-1 ligand or receptor (and show little to no *daf-7* expression in their ASJ neurons on *E. coli*, see Figure 1B) were capable of upregulating *daf-7* expression in ASJ upon exposure to *P. aeruginosa* (Figure 1—Figure Supplement 1). Given the previously established function of *daf-7* expression in the ASJ neurons of hermaphrodites in promoting pathogen avoidance behavior (Meisel et al., 2014), these results predict that mutants in the PDF-1 pathway should have no defects in their ability to avoid a lawn of pathogenic *P. aeruginosa*. As expected, we observed that while *daf-7* mutant hermaphrodites fail to avoid a lawn of pathogenic bacteria, the *pdf-1* and *pdfr-1* mutant hermaphrodites appear wild-type for their ability to perform this behavior (Figure 1E). Together this data suggests that the PDF-1 signaling pathway acts both cell-specifically—on the ASJ neurons—as well as sex-specifically— in males—to regulate *daf-7* expression and its effects on downstream behavioral programs.

### PDF-1 signals to the ASJ neurons to promote *daf-7* expression and mate-searching behavior

The PDF-1 neuropeptide ligand is secreted from multiple neurons in the head region of the animal where a similarly large number of neurons express the PDFR-1 receptor (Barrios et al., 2012; Janssen et al., 2009; Meelkop et al., 2012). To identify the relevant site of action for this pathway in the regulation of *daf-7* expression in males, we used the *pdfr-1(ok3425)* mutant animals and introduced *pdfr-1* cDNA transgenes into specific neurons using heterologous cell-specific promoters. We observe that while the *pdfr-1* mutant males lack significant *daf-7* expression in the ASJ neurons, introduction of a genomic DNA fragment carrying the *pdfr-1* locus could fully rescue this phenotype and restore *daf-7* expression in the ASJ neurons (Figures 2A and B). We next asked if *pdfr-1* expression in the ASJ neurons alone was sufficient to rescue the mutant phenotype of the *pdfr-1(ok3425)* males and drove the expression of the B isoform of *pdfr-1* under the control of the ASJ-specific *trx-1* promoter. We observe that expression of *pdfr-1* in this single-pair of neurons (ASJ) was indeed sufficient to rescue *daf-7* expression into the ASJ neurons of the mutant male animals, suggesting that this pathway signals to ASJ directly to influence *daf-7* expression specifically in the male (Figure 2A and B).

**Figure 2.**
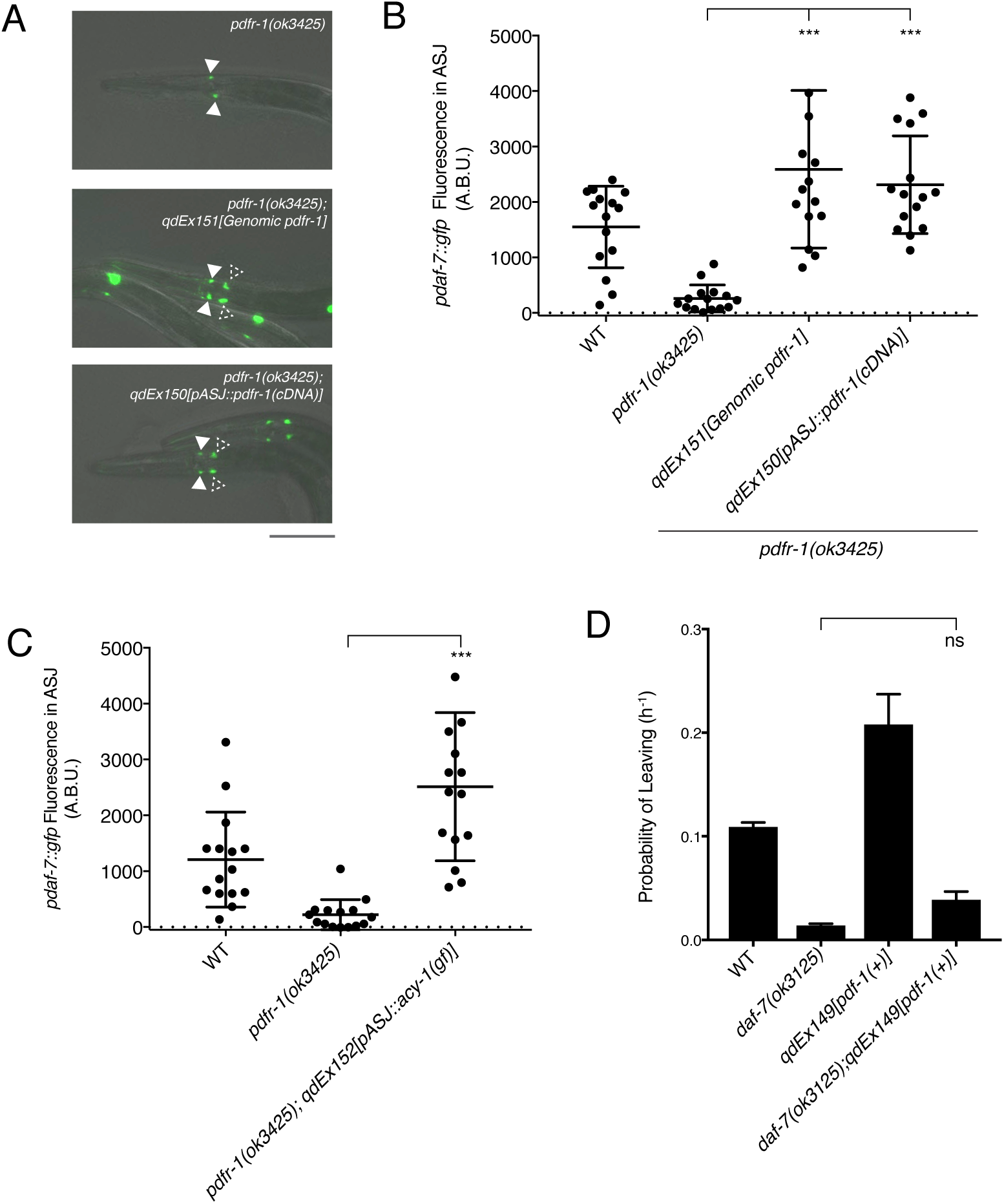
PDF-1 signaling functions in ASJ to regulate *daf-7* expression and mate-searching behavior in male *C. elegans*. (A) *pdaf-7::gfp* expression in *pdfr-1(ok3425)* mutant (top), genomic rescue (middle), and ASJ-specific rescue (bottom) male animals. Filled arrowheads indicate the ASI neurons; dashed arrowheads indicate the ASJ neurons. Scale bar indicates 50 μm. (B) Maximum fluorescence values of *pdaf-7::gfp* in the ASJ neurons of *pdfr-1* rescue males. ***p< 0.001 as determined by ordinary one-way ANOVA followed by Dunnett’s multiple comparisons test. Error bars represent SD. n=15 animals for all genotypes. (C) Maximum fluorescence values of *pdaf-7::gfp* in the ASJ neurons of males expressing the gain-of-function ACY-1(P260S) cDNA specifically in ASJ. ***p< 0.001 as determined by unpaired t-test with Welch’s correction. Error bars represent SD. n=15 animals for all genotypes. (D) Probability of leaving values for epistasis experiment between *daf-7(ok3125)* and a PDF-1 overexpressing line. Values plotted are the mean + SEM for three independent experiments. Significance determined by unpaired t-test with Welch’s correction. ns, not significant. n=60 total animals for all genotypes except the *daf-7(ok3125); qdEx149* strain where n=48.

The PDFR-1 receptor is a secretin-family G-protein coupled receptor (GPCR), which has been shown to stimulate G*α*s signaling and upregulation of cAMP production in transfected cells as well as in both *Drosophila melanogaster* and *C. elegans* neurons (Figure 1A; Flavell et al., 2013; Hyun et al., 2005; Janssen et al., 2008; Lear et al., 2005; Mertens et al., 2005; Shafer et al., 2008). Using a gain-of-function variant of the adenylate cyclase, ACY-1 (Flavell et al., 2013; Saifee et al., 2011; Schade et al., 2005), we asked if activation of the pathway downstream of PDFR-1 specifically in ASJ was sufficient to rescue the defects in *daf-7* expression that we observe in the *pdfr-1* mutant males. We observe that in *pdfr-1* mutant males with transgenic expression of the *acy-1(gf)* cDNA only in the ASJ neurons, *daf-7* expression was fully rescued (Figure 2C). This ability to bypass PDFR-1 by activation of cAMP production specifically in the ASJ neuron pair further suggest that the PDF-1 signaling pathway acts directly on the ASJ neurons in order to regulate *daf-7* expression in male animals.

Expression of *daf-7* in the ASJ neuron pair of males is required for the male-specific mate-searching behavioral program (Hilbert and Kim, 2017), while the PDF-1 pathway has similarly been implicated as a regulator of this same behavior (Barrios et al., 2012). Given the role that this PDF-1 pathway plays in regulating the expression of *daf-7*, we set out to determine if the effects of the PDF-1 pathway on mate-searching behavior are the result of PDF-1 and DAF-7 functioning through a single pathway or through separate parallel pathways. It has been previously established that overexpression of the *pdf-1* genomic sequence confers increased mate-searching behavior in male animals (Figure 2D; Barrios et al., 2012). We took advantage of this observation to perform epistasis between the PDF-1 and DAF-7 pathways by introducing a *daf-7* mutation into these transgenic PDF-1 overexpressing lines. Importantly, whereas the males carrying our overexpression construct alone display increased leaving rates as previously reported, this effect was entirely suppressed by mutation of *daf-7* (Figure 2D and Figure 2— Figure Supplement 1). These data suggest that the PDF-1 pathway regulates this male-specific behavior through the regulation of *daf-7* expression in the ASJ neurons of adult male animals, although our data do not exclude the possibility that PDF-1 may also regulate mate-searching through additional parallel pathways.

### Sex differences in PDF-1 receptor expression underlie the sex-specific regulation of *daf-7* transcription in ASJ

The sex-specificity of the effects of the PDF-1 pathway on *daf-7* regulation and mate searching behavior is intriguing given very little evidence of differences in the expression or function of this neuropeptide pathway between the two *C. elegans* sexes (Barrios et al., 2012; Janssen et al., 2008; 2009). It was recently shown that *pdf-1* is produced by the newly identified male-specific MCM neurons and is required for the regulation of sex-specific learning in males, but interestingly, ablation of these neurons has no effect on mate-searching behavior (Sammut et al., 2015). Nevertheless, we wondered if there might be unidentified sex differences in the signaling or expression of this PDF-1 neuropeptide pathway in neurons such as ASJ, which would confer sex-specific activity of the PDF-1 pathway on the regulation of *daf-7* gene expression. To this end, we asked if activation of the PDF-1 signaling pathway in the ASJ neurons of hermaphrodites might be sufficient to drive *daf-7* expression inappropriately in these animals. We first looked at hermaphrodite animals carrying the same ASJ-expressed *acy-1(gf)* transgene and observed significant upregulation of *daf-7* expression in the ASJ neurons of these hermaphrodites (Figure 3A). We next asked whether we could observe *daf-7* expression in the ASJ neurons of hermaphrodite animals with heterologous expression of *pdfr-1* in only the ASJ neurons. Strikingly, we found that in hermaphrodites with overexpression of *pdfr-1* cDNA in the ASJ neurons, *daf-7* expression is also upregulated in the ASJ neurons similar to what we observed in the *acy-1(gf)* transgenic strains (Figure 3B). We also quantified *daf-7* expression in the ASJ neurons of hermaphrodites carrying the genomic *pdfr-1* fragment with all the endogenous regulatory sequence and observed no upregulation of expression in those animals. This control suggests that *daf-7* expression in ASJ cannot be triggered simply as the result of overexpression of *pdfr-1* (Figure 3B), rather, these results suggest that expression of PDFR-1 specifically in the hermaphrodite ASJ neurons is sufficient to allow *daf-7* expression in these neurons.

**Figure 3.**
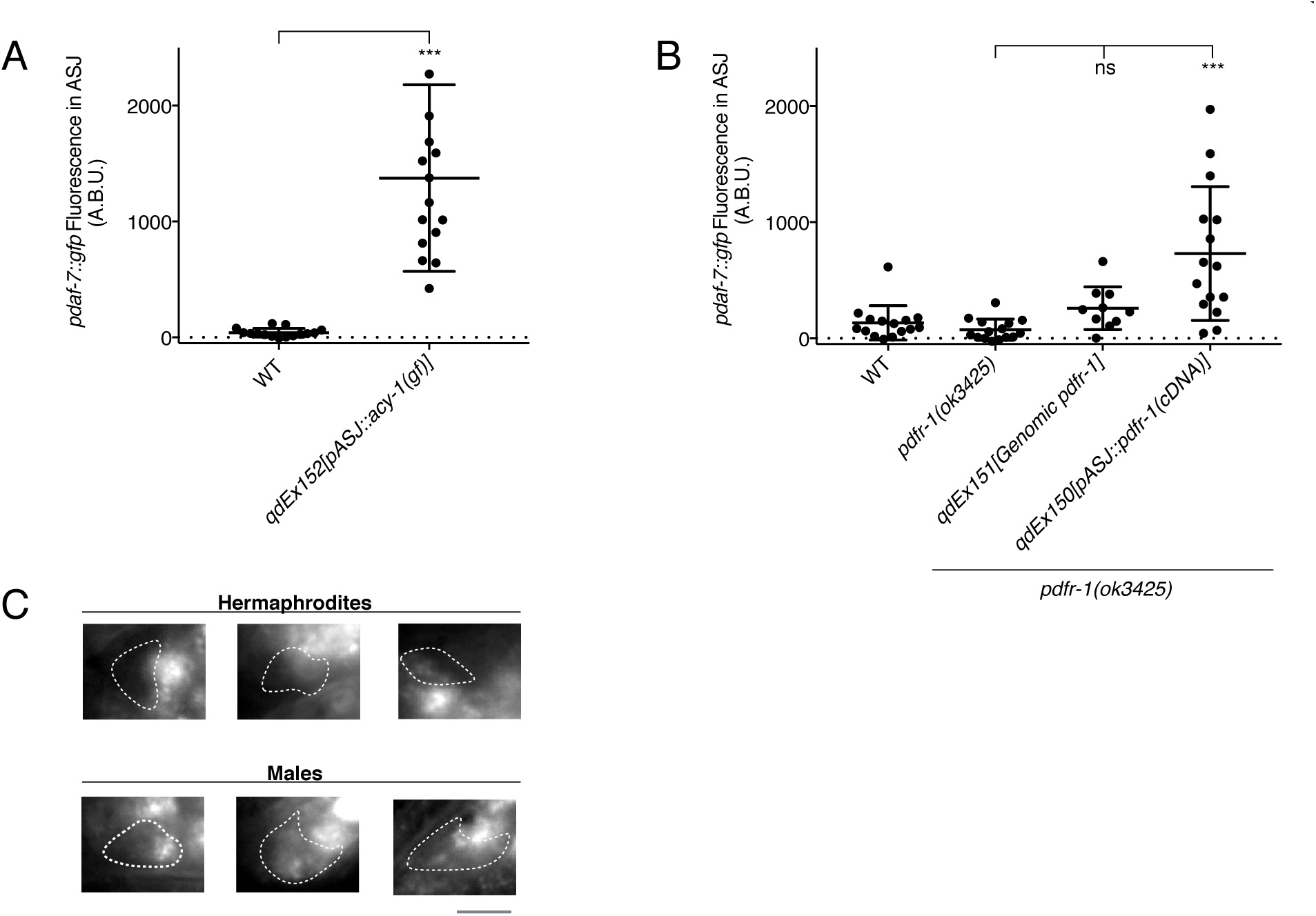
Heterologous activation of the PDF-1 pathway in ASJ is sufficient to activate *daf-7* transcription in adult hermaphrodites. (A) Maximum fluorescence values of *pdaf-7::gfp* in the ASJ neurons of WT hermaphrodites and hermaphrodites where ACY-1 has been activated (via the gain of function P260S mutant) specifically in ASJ. ***p< 0.001 as determined by unpaired t-test with Welch’s correction. Error bars represent SD. n=15 animals for both genotypes. (B) Maximum fluorescence values of *pdaf-7::gfp* in the ASJ neurons of hermaphrodites overexpressing *pdfr-1* from either a genomic fragment or under the control of a heterologous ASJ-specific promoter. ***p< 0.001 as determined by ordinary one-way ANOVA followed by Dunnett’s multiple comparisons test. Error bars represent SD. ns, not significant. n=15 animals for all genotypes except *pdfr-1(ok3425); qdEx151* where n=10 animals. (C) Representative FISH images of endogenous *pdfr-1* mRNA molecules in adult hermaphrodite (top) and male (bottom) ASJ neurons. White dotted lines indicate outline of ASJ cell body as localized by *ptrx-1::gfp*. Animals imaged carried a transgene with the *pdfr-*1 genomic locus to increase expression (see Figure 3—Figure Supplement 1). Scale bar represents 5 µm.

Our results suggest a possible hypothesis that expression differences at the level of the PDFR-1 receptor might explain the sex-specific regulation of this pathway on *daf-7* expression in the ASJ neurons. To further assess this possibility, we used fluorescence *in situ* hybridization (FISH) to examine the transcription of *pdfr-1* in male and hermaphrodite animals. We generated fluorescent probes for a region of the *pdfr-1* coding sequence that is shared among all isoforms and verified the specificity of these probes by examining expression in the *pdfr-1(ok3425)* deletion mutant, where we observed no fluorescent signal (Figure 3—Figure Supplement 1B). Similarly, when we looked at *pdfr-1* mRNA transcripts in our ASJ-specific rescue lines, we could only observe fluorescence in the ASJ neurons (Figure 3—Figure Supplement 1D), suggesting that these probes are highly specific for the *pdfr-1* coding sequence. Imaging of *pdfr-1* transcription in WT animals revealed a diffuse expression pattern with fluorescent signal observable in muscle tissue as well as in neurons, but with few cells having strong signal and many cells with only scattered fluorescent spots, including ASJ (Figure 3—Figure Supplement 1A). To corroborate and confirm these observations, we also imaged *pdfr-1* transcripts in animals carrying our genomic rescuing fragment, which amplified probe fluorescence throughout the nervous system and muscle (Figure 3—Figure Supplement 1C). We expect that because of the intact endogenous regulatory sequence on this genomic fragment, the mRNA localization we observe in this strain should still be representative of the wild-type expression pattern of *pdfr-1*. Using these animals, we observe more abundant *pdfr-1* mRNA in the ASJ neurons of a number of adult male animals (Figure 3C). In contrast, we did not observe *pdfr-1* mRNA in the ASJ neurons of hermaphrodite animals even in this overexpression context, suggestive that this gene is expressed in a sexually dimorphic manner in the ASJ neurons (Figure 3C).

### The PDF-1-DAF-7 neuroendocrine signaling cascade regulates sex-specific behavior through the sex-shared ASJ neurons

Building on our previous work on the sexually dimorphic regulation of the neuroendocrine gene *daf-7* and its role in promoting male decision-making behaviors (Hilbert and Kim, 2017), we have presented here a set of experiments which implicate the PDF-1 neuropeptide signaling pathway as a critical male-specific regulator of *daf-7* expression in the ASJ neurons. Our data suggest that sexual dimorphism in the expression of the PDF-1 receptor, PDFR-1, may serve as a gating mechanism, allowing the ASJ neurons of adult male *C. elegans* to respond to the PDF-1 ligand. We suggest that this ligand-receptor interaction activates a downstream signaling cascade in ASJ terminating in the transcriptional activation of *daf-7*, which in turn promotes male-specific decision-making behaviors (Figure 4, left). We hypothesize that the relative lack of expression of *pdfr-1* in the hermaphrodite ASJ neurons prevents the activation of this pathway and consequently *daf-7* expression is not induced under normal growth conditions in adult hermaphrodites (Figure 4, right). Strikingly, heterologous expression of the PDF-1 receptor in the hermaphrodite ASJ neurons was sufficient to drive *daf-7* expression in an inappropriate physiological context (the hermaphrodite nervous system), suggesting the integral role of this pathway in facilitating sex-specific differences in gene expression and behavior.

**Figure 4.**
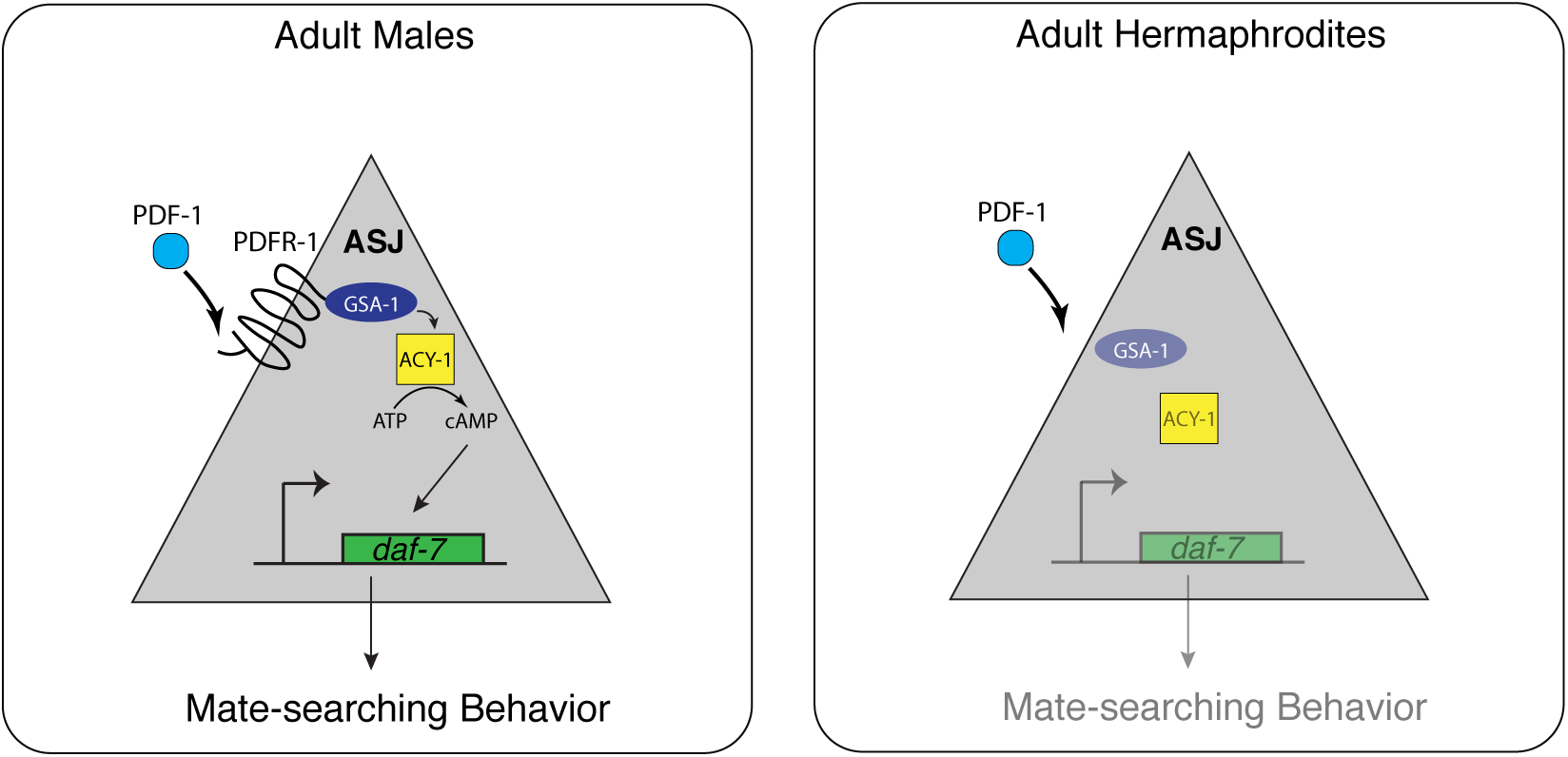
A sexually dimorphic neuroendocrine cascade functions in the sex-shared nervous system to regulate decision-making behaviors. We show that the PDF-1 neuropeptide pathway functions in facilitating the sex-specific expression of the TGF*β* ligand, DAF-7, in the ASJ chemosensory neurons and through this regulation has downstream effects on male-specific exploratory behaviors. Sex differences at the level of the expression of the PDFR-1 receptor enable a gating mechanism on the ability of these sex-shared neurons to respond to an endogenous neuroendocrine cue and provide insight into how shared neuronal circuitry can achieve sex-specific differences in gene expression and behavior.

While recent work has revealed sexual dimorphisms at the level of gene expression, neuronal connectivity and neurotransmitter release in the sex-shared nervous system of *C. elegans* (Hart and Hobert, 2018; Hilbert and Kim, 2017; Oren-Suissa et al., 2016; Pereira et al., 2015; Ryan et al., 2014; Serrano-Saiz et al., 2017a; 2017b; Weinberg et al., 2018), the role of neuromodulators and other neuroendocrine signals in facilitating sex-specific responses of neurons in the shared neuronal circuitry has been relatively unexplored. Here, we propose a model in which two pathways, the PDF-1 and DAF-7/TGF*β* pathways, act in concert as a neuroendocrine signaling cascade to regulate sex-specific behavior within the context of the sex-shared ASJ neurons (Figure 4). Our data suggest that the PDF-1 pathway functions in tuning the response of the ASJ neurons to this endogenous neuromodulator in a sex-specific manner via differential expression of PDFR-1. Interestingly, recent work in mice has uncovered a similar phenomenon wherein the neuromodulator oxytocin facilitates sex-specific social preference in male mice by modulating the ability of subsets of neurons to respond to social cues (Yao et al., 2017). The parallels between this work and ours underscore the role of neuroendocrine signaling through sex-shared nervous system components in shaping sexually dimorphic neuronal activity and behavior in evolutionarily diverse animals.

## MATERIALS AND METHODS

### *C. elegans* Strains

*C. elegans* strains were cultured as previously described (Brenner, 1974; Hilbert and Kim, 2017). For a complete list of strains used in this study please see Supplementary File 1.

### Cloning and Transgenic Strain Generation

For the *pdf-1* overexpression transgene, a 6.5 kb region of sequence containing the *pdf-1* promoter, coding sequence and 3’UTR were amplified from the fosmid WRM0641dA07 from the Moerman fosmid library. This fragment was cloned into the pUC19 vector backbone by Gibson assembly (Gibson et al., 2009).

For the ASJ-specific *pdfr-1* rescue construct, the B isoform of the *pdfr-1* cDNA with no stop codon was amplified from cDNA generated with an Ambion RetroScript kit using primers based on previously described annotation of the isoform (Barrios et al., 2012). The *trx-1* ASJ-specific promoter was amplified as previously described (Hilbert and Kim, 2017). An F2A::mCherry fragment was amplified off a plasmid that was a gift from C. Pender and H.R. Horvitz. All fragments were cloned into the pPD95.75 backbone with an intact *unc-54* 3’UTR by Gibson assembly. Genomic rescue of *pdfr-1* was done by injection of the WRM0629dH07 fosmid from the Moerman fosmid library.

For the ACY-1(gf) construct, the 3.8 kb *acy-1(P260S)* fragment was amplified from genomic DNA extracted from the strain CX15050 (gift from S. Flavell and C. Bargmann) which carries a transgenic array with the *acy-1(P260S)* cDNA under the control of a different promoter. This fragment was cloned into a plasmid backbone carrying the *trx-1* promoter and *unc-54 3’UTR.* All fosmids and plasmids were verified by sequencing and injected at a concentration of 50 ng/μL along with a plasmid carrying *pofm-1::gfp* at 50 ng/μL as a co-injection marker. At least 3 independent transgenic lines were obtained and analyzed for each construct and one representative line is shown. For a list of all primers used in this paper, please see Supplementary File 2.

### Measurement of Gene Expression in ASI and ASJ neurons

Quantification of *daf-7* expression was performed as described in (Hilbert and Kim, 2017). All adult quantifications were done on animals 72 hours after egg lay. Quantification of animals on

*P. aeruginosa* were performed as before.

### Mate-Searching Assays

Mate-searching assays were performed as previously described (Hilbert and Kim, 2017; Lipton et al., 2004).

### Fluorescence In Situ Hybridization

FISH was performed as previously described (Hilbert and Kim, 2017; Raj et al., 2008). The *pdfr-1* probe was constructed by pooling together 36 unique 20 nucleotide oligos that tile across base-pairs 580-1540 in the *pdfr-1* B-isoform cDNA. This sequence is contained in all isoforms of *pdfr-1* so should anneal to any endogenous *pdfr-1* mRNA. After pooling, oligos were coupled to Cy5 and then purified by HPLC.

### *P. aeruginosa* Lawn Avoidance Assays

*P. aeruginosa* plates were prepared as described in (Hilbert and Kim, 2017). Animals were synchronized by treatment with bleach and allowed to hatch and arrest as L1 larvae before being dropped onto *E. coli* plates. L4 animals were transferred to the center of the *P. aeruginosa* lawn, incubated at 25°C and then scored for avoidance after 16 hours.

### Statistical Analysis

All statistical analysis was performed using the Graphpad Prism software. Statistical tests used for each experiment are listed in the figure legend.

## ACKNOWLEDGEMENTS

We thank C. Bargmann, S. Flavell, H.R. Horvitz, J. Kaplan, and the *Caenorhabditis* Genetics Center (which is funded by the NIH Office of Research Infrastructure Programs P40 OD010440) for strains and reagents. We thank S. Flavell and members of the Kim and Horvitz labs for helpful discussions in the development of the manuscript.

## Supplementary File 1. *C. elegans* strains used in this study

A comprehensive list of the strains used in this study. Strain source (this study or others) is indicated.

## Supplementary File 2. Oligos used for transgenic strain construction and *pdfr-1* FISH

A comprehensive list of the oligos created for this study. All sequences are listed in the 5’ to 3’ direction.

**Figure 1—Figure Supplement 1.**
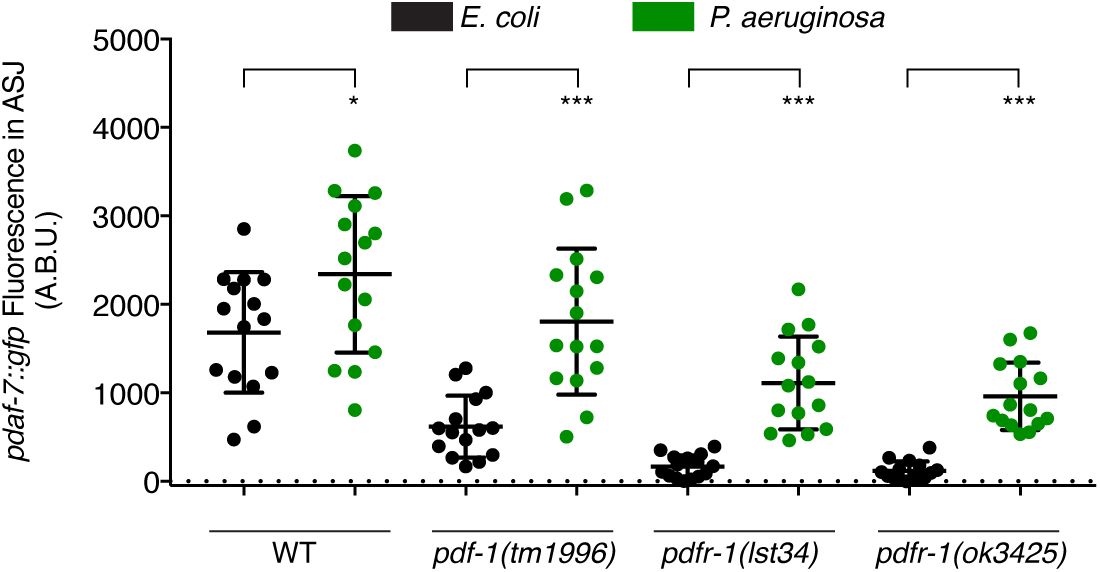
PDF-1 pathway mutant males can upregulate *daf-7* expression in ASJ in response to *P. aeruginosa* exposure. Maximum fluorescence values of *pdaf-7::gfp* expression in ASJ for males exposed to *E. coli* (black) or *P. aeruginosa* (green) for 16 hours. ***p< 0.001, *p< 0.05 as determined by unpaired t-tests with Welch’s correction. n=15 animals for all genotypes and conditions.

**Figure 2—Figure Supplement 1.**
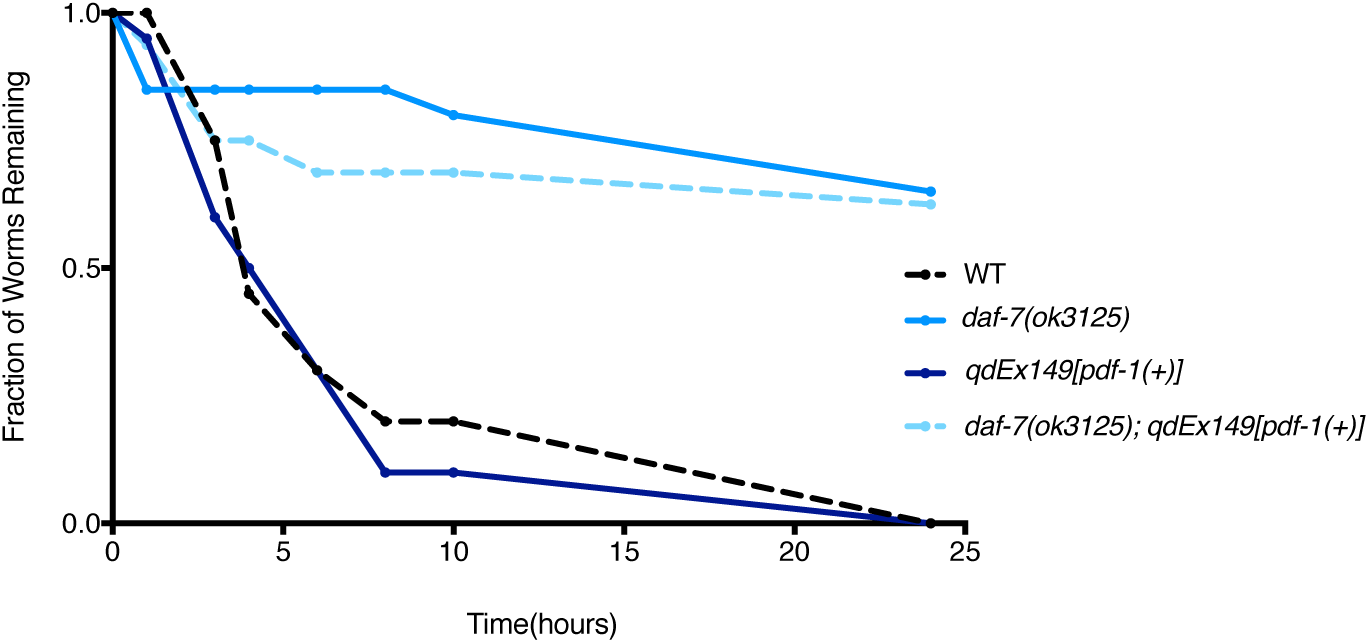
The PDF-1 pathway regulates mate-searching behavior via male-specific modulation of *daf-7* expression in ASJ. Representative mate-searching data for epistasis experiment between PDF-1 and DAF-7. Leaving curves are shown for WT (dashed black), *daf-7* mutant (light blue), *pdf-1(*+) transgenics (dark blue), and the *daf-7; pdf-1(*+) double (light blue dashed) males. n=20 animals for each curve shown except for *daf-7;pdf-1(*+) where n=16 animals.

**Figure 3—Figure Supplement 1.**
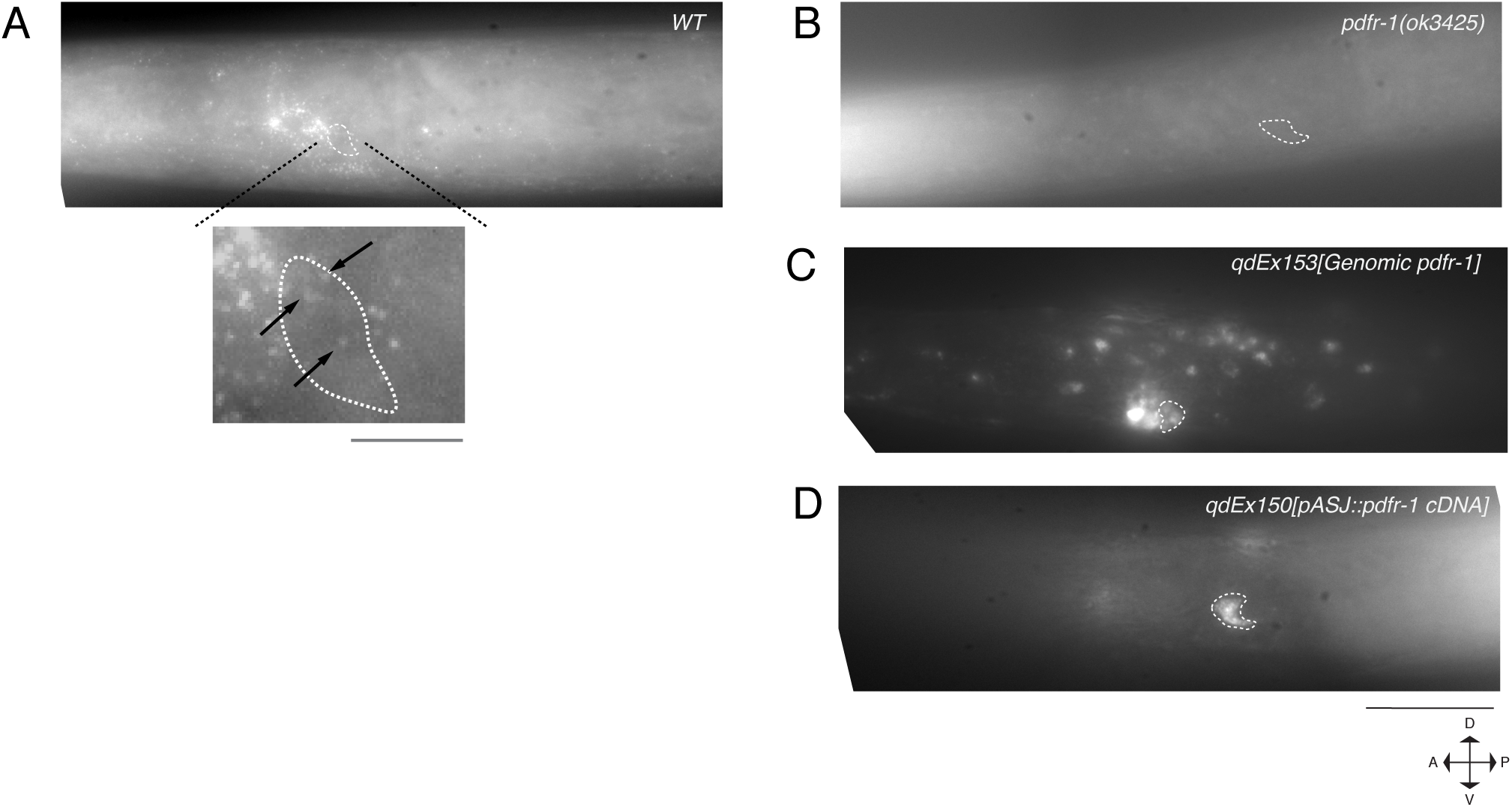
FISH imaging of endogenous *pdfr-1* mRNA transcripts. (A) *pdfr-1* transcripts in a WT male animal. mRNA can be observed in neurons as well as diffusely in muscle and throughout the body. Magnified image of an ASJ neuron is shown on bottom. Arrows indicate *pdfr-1* transcripts. Scale bar for magnified image represents 5 µm. (B) *pdfr-1* mRNA in a *pdfr-1(ok3425)* mutant male. No mRNA can be visualized indicating the specificity of the designed probe set. (C) *pdfr-1* transcripts in a male carrying a genomic fragment including the *pdfr-1* locus on an extrachromosomal transgenic array. Animals carrying this array have increased *pdfr-1* expression but with all the endogenous regulatory sequence, facilitating easier analysis of expression in specific neurons, such as ASJ. (D) *pdfr-1* mRNA in a *pdfr-1(ok3425)* male with heterologous expression of *pdfr-1* cDNA in the ASJ neurons. No transcripts can be observed outside of the ASJ neurons. All images presented are maximum intensity z-projections of 28 stacked exposures of *pdfr-1* mRNA. ASJ neurons are outlined in each image by a white dotted line as localized by either *ptrx-1::gfp* or *pdaf-7::gfp* (for the ASJ rescue line). Animals are representative samples from mixed stage populations. All images were taken at 100x magnification. Scale bar indicates 50 µm.

